# Insulin resistance-driven beta-cell adaptation in mice: Mechanistic characterization and 3D analysis

**DOI:** 10.1101/2023.01.09.523222

**Authors:** Alexandrine Liboz, Carine Beaupere, Natacha Roblot, Jean-Yves Tinevez, Sandra Guilmeau, Anne-Françoise Burnol, Dalale Gueddouri, Xavier Prieur, Bruno Fève, Ghislaine Guillemain, Bertrand Blondeau

## Abstract

**Aims/hypothesis:** Pancreatic beta cells secrete insulin to control glucose homeostasis. Beta cells can also adapt their function and mass when more insulin is required, especially in situations of insulin resistance (IR). Beta-cell mass adaptation can be achieved through either beta-cell proliferation or beta-cell neogenesis, a process that involves de novo beta-cell production from precursor cells. Signals and mechanisms that control adult beta-cell neogenesis and regulate the balance between beta-cell proliferation and/or beta-cell neogenesis still need to be fully deciphered. To do so, we previously developed a mouse model of pancreatic adaptation in response to a severe insulin resistance induced by a chronic glucocorticoid (GC) treatment. We observed a massive insulin production due to beta-cell adaptation by both proliferation and neogenesis. In the present study, we aimed at further characterizing beta-cell adaptation in response to mild or severe IR by studying various GC doses, along with other pharmacological or genetic models of IR. Further, we characterized the impact of aging on pancreatic adaptation in response to GC-induced IR. Finally, we precisely quantified adult beta-cell neogenesis by developing an original 3D method of beta-cell mass analysis in toto after organ clearing.

**Methods:** Glucose metabolism, insulin secretion and pancreatic beta-cell adaptation were studied in mice rendered IR either by adipose tissue specific invalidation of SEIPIN, by chronic treatment with the insulin receptor antagonist S961 or by chronic treatment with several doses of GC both in young and aged mice. Moreover, we developed and used an unbiased-3D analysis of beta cells on whole cleared pancreas.

**Results:** We demonstrated that beta-cell neogenesis - reflected by an increase in islet density - is constantly observed in response to genetically- or pharmacology-induced (S961 or GC) IR. Next, we observed that pancreatic adaptation mechanisms are closely defined by the level of IR. Indeed, mild IR induced by low dose of GC resulted in functional adaptation solely, while more severe IR induced by higher doses of GC resulted in an increase in both islet density and mean islet size, reflecting beta-cell neogenesis and proliferation, respectively. Then, we showed that in older mice, beta-cell adaptation through neogenesis is preserved in response to IR. Finally, using a new and unbiased 3D analysis, we confirmed the increase in islet density and mean islet size after GC’s treatment.

**Conclusions/interpretation:** Our results present evidence that beta-cell neogenesis is a preferential mechanism of pancreatic adaptation to increase insulin secretion in response to IR in mice. Moreover, aging does not preclude beta-cell neogenesis, suggesting that it could be triggered in elderly to compensate for IR. Finally, our innovative technique of 3D analysis of whole pancreas confirms the existence of adult beta-cell neogenesis and offers a new avenue to study islet cells and pancreas adaptation.

**Research in context:** *What is already known about this subject?:*

- Insulin resistance can be compensated by improved insulin secretion and increased beta-cell mass.
- New beta cells can be formed in the pancreas of adult mice through the differentiation of precursors, a process known as neogenesis.
- We previously demonstrated that glucocorticoid (GC) -induced insulin resistance leads to enhanced beta-cell proliferation and neogenesis.

*What is the key question?:*

- Is adaptive beta-cell neogenesis specific to GC-induced insulin resistance and persists in old mice ?

*What are the new findings?:*

- Insulin resistance, either genetically- or drug-induced, is a key driver to induce adaptive beta-cell neogenesis in the mouse pancreas.
- Aging does not prevent the induction of beta-cell neogenesis in response to insulin resistance.
- Three-dimension analysis on cleared pancreas confirms beta-cell neogenesis in mouse models of GC-induced insulin resistance.

*How might this impact on clinical practice in the foreseeable future?:*

- The mouse model of adaptive beta-cell neogenesis will be helpful to define new therapeutic targets to induce the formation of new beta cells and treat diabetes.

## Introduction

Pancreatic beta cells that secrete insulin - the only hypoglycemic hormone of the body - play a central role in the regulation of blood glucose. Insulin both activates glucose storage in fat, liver and skeletal muscles and inhibits glucose production by the liver, allowing a fine regulation of blood glucose levels. Moreover, beta cells are able to increase their mass and insulin production when required, to adapt to the increased demand in insulin induced by hyperglycemia and insulin resistance (1). Because of their central role, beta-cell loss and/or dysfunction are involved in the development of diabetes, which are characterized by a deficiency in insulin secretion caused by either the total or partial loss of beta-cell mass, in type 1 and type 2 diabetes respectively (2).

To compensate for the loss of beta cells in diabetes, research strategies have been explored but dampened by our limited understanding of the mechanisms underlying regeneration of pancreatic beta cells in the adult pancreas. Models of pancreatectomy and pancreatic duct ligation have been studied in mice and rats, and showed an increase in beta-cell mass in the adult rodent pancreas through the formation of new beta cells, a process known as neogenesis (11). More specifically, such neogenesis required the re-expression of a transcription factor involved in the fetal development of the endocrine pancreas, Neurogenin3 (*NGN3*). In order to characterize the origin of these *neo* formed beta cells, cell lineage studies following the expression of the transcription factor *SOX9* were performed in animal models and found precursor cells originating from pancreatic ducts (12). However, other groups were unable to reproduce these results and there is a current controversy about the origin of the newly formed beta cells (13, 14).

In Humans, beta cells neogenesis was suggested by the group of Butler that demonstrated an increase in beta-cell mass to respond to a physiological insulin resistance induced by pregnancy through an increase in the number of islets per pancreas, without proliferation of pre-existing beta cells (15). Similarly, the work by Mezza and colleagues showed that non-pathological insulin resistance is associated with pancreatic adaptation by neogenesis in non-diabetic individuals (16). These two studies therefore showed that in response to insulin resistance, the human pancreas was able to adapt by triggering the formation of new beta cells.

Triggering beta-cell neogenesis represents promising therapeutic avenues for beta-cell regeneration to treat insulin deficiency. Yet, it is not known if beta cell neogenesis is a main adaptive mechanism in response to insulin resistance as compared with other pancreatic adaptation mechanisms (i.e. increase in beta-cell function or proliferation). To gain strong insights on beta-cell neogenesis *in vivo*, the use of animal models of insulin resistance either physiological, genetic or drug-induced is required. Several drugs can induce or mimic insulin resistance, such as a long-term treatment with glucocorticoids (GC) (17) or with S961, an insulin receptor antagonist causing hyperinsulinemia and hyperglycemia (18). GC are known to alter glucose homeostasis, both by reducing insulin sensitivity at the liver, muscle and adipose tissue levels, and by modulating insulin resistance (19, 20, 21, 22). Previously, we developed a model of GC-induced insulin resistance using chronic corticosterone (CORT) treatment and characterized beta-cell adaptation in mice (17). We showed a massive increase in beta-cell mass due to an augmented cell proliferation and an increase in islet density, suggesting beta-cell neogenesis. Accordingly, we observed an increase in *NGN3* expression along with other genes involved in beta-cell neogenesis. Interestingly, more islets were connected to or in vicinity of ducts, although cell lineage revealed that newly formed beta cells did not derive from *SOX9* nor *NGN3* expressing cells. Moreover, beta-cell neogenesis in our model achieved partial regeneration even after beta-cell depletion with streptozotocin (STZ). Finally, we demonstrated that beta-cell neogenesis was not a direct effect of GC, but rather due to the presence of pro-neogenic factors in the serum of GC-treated mice.

Our model proved the possibility of a massive increase in beta-cell mass in the pancreas of young adult mice and involving neogenesis. The aim of our present study is to better characterize pancreatic adaptation in response to GC-induced insulin resistance, i.e. its dynamic and its specificity compared to other model of insulin resistance. Indeed, severe insulin resistance can also be achieved using the S961 - an insulin receptor antagonist – who was reported to induce pancreatic adaptation by beta-cell proliferation in both rats and mice (23, 24). However, beta-cell neogenesis has not been investigated in this model. Apart from pharmacological models of insulin resistance, genetic-lipodystrophies, leading severe insulin resistance could be studied (25). Among them, patients with a Berardinelli-Seip syndrome, carrying mutations in the *BSCL2* gene - encoding seipin, a protein required in lipid droplets formation and maintenance within adipocytes - display a near complete loss of adipose tissue causing a severe insulin resistance and ultimately diabetes. In mice, *BSCL2* gene deletion leads to nearly complete lipodystrophy associated with severe insulin resistance (25). In addition, adipocyte-specific deletion of *BSCL2* gene in mice was developed and leads to progressive lipodystrophy ultimately associated with glucose intolerance and insulin resistance (26).. Interestingly, pancreatic beta cells function has been assessed in the constitutive model of *BSCL2* deletion but pancreatic cell adaptation remains to be assessed in adipose-specific BSCL2-deleted mice (27).

To further characterize the relationship between GC-induced insulin resistance and beta-cell adaptation, we focused on the level of insulin resistance needed to induce neogenesis by treating mice with different doses of GC. Because in humans insulin resistance often appears in elderlies, we also asked whether beta-cell adaptation to GC-treatment could also be observed in old mice and whether similar mechanisms are at play in-between young and aged mice. Indeed, several studies demonstrated a decreased capacity in beta-cell proliferation and islet regeneration with age, although neogenesis itself in aged mice was not assessed (28).

Finally, although we and others demonstrated a clear increase in pancreatic islets density (notably small islets) by 2D analysis techniques on tissue sections, the existence of adult beta-cell neogenesis remains controversial nowadays. To ascertain its existence, we aimed at developing a protocol for the clearing and three-dimensional analysis of the whole endocrine pancreas from control (VEH) and CORT-treated, young and old mice, to precisely quantify and characterize islets throughout the whole pancreas.

In this study, we showed in mice that 1) treatments with the insulin receptor antagonist S961 or a genetic model of lipodystrophy induce insulin resistance and lead to pancreatic adaptation through beta-cell neogenesis; 2) beta-cell neogenesis persists as a main adaptive mechanism in old mice; 3) there is a threshold of GC treatment and insulin resistance necessary to trigger adaptation of beta-cell mass and function; 4) beta-cell neogenesis can be confirmed and precisely quantified by an innovative 3D analysis of cleared adult mice pancreas, demonstrating that regenerative therapies directly targeting beta-cell neogenesis could be envisioned in the near future.

## Research Design and Methods

### Animals

All procedures involving experimental animals were performed in accordance with the principles and guidelines established by INSERM and were approved by the local animal care and use committee (Charles Darwin ethic committee, Paris, France). C57BL/6J male mice were obtained from Charles River Laboratories (Saint-Germain-sur-l’Arbresle, France) at the age of 8 weeks or 52 weeks. Mice were chow fed *ad libitum* and housed in 12-h light/dark cycles.

### Bscl2^lox/lox^ × Adipoq-CreER mice

Bscl2^lox/lox^ mice were crossed with Adipoq-CreER (29) mice to generate Bscl2^lox/lox^ X Adipoq-CreER mice as previously described (26). Cre activation was performed by 5 consecutive days of intraperitoneal Tamoxifen (Tam) injection resuspended in sunflower oil:ethanol (9:1) at 8 weeks-old, and then one injection every month until 6 months. Bscl2^lox/lox^ X Adipoq-CreER mice that receive Tam will be referred as Adipo Seipin KO and Bscl2^lox/lox^ mice treated with Tam will be referred as Adipo Seipin^lox/lox^. Insulin tolerance test were performed after 6-hours fasting as previously described (26). All mice (males, n=8 to 11 per group) were euthanized in the random fed state between 8 a.m and 10 a.m. The ethics committee of the French national veterinary agency approved all animal protocols used in this study.

### Chemicals

#### Corticosterone (CORT)

Animals were treated with either CORT dissolved in ethanol (100, 50 or 25 μg/ml, as indicated) (Sigma-Aldrich, St. Louis, MO) or vehicle (VEH) (1% ethanol) in drinking water for 3 to 4 weeks.

#### S961

ALZET® mini osmotic pumps were implanted subcutaneously in eight-week-old male C57Bl/6 J mice to infuse either S961, an insulin receptor (IR) peptide antagonist (Novo Nordisk) (20 nmol/week), or PBS, as VEH control, at 0.5 μl/h for 1 week as previously described (30).

### Insulin Tolerance Tests (ITT)

Insulin tolerance test (ITT) were performed as previously described (31). Briefly, after a 6-h fast, insulin (1 unit/kg of body weight) was injected intraperitoneally. Blood glucose levels were measured before and 15, 30, 60, and 120 min after the injection, using a glucometer (Accu-checkPerforma, Roche). Slope of early insulin sensitivity were calculated as the difference of blood glucose levels 15 minutes after insulin and before insulin injection and divided by time (15 minutes). Slopes were expressed as mg/dl/min.

### Hormonal Assays

Blood was collected by intracardiac puncture. Serum was collected after 5 min of centrifugation at 2000g at 4°C. Insulin levels were measured using mouse insulin immunoassay (Merck, Millipore) according to manufacturer’s instructions.

### Islet Isolation

Mouse islets were isolated after injecting a collagenase solution (1 mg/mL) (Sigma-Aldrich) in the bile duct and handpicked under a binocular microscope (Leica Microsystems, Wetzlar, Germany) as described in (32).

### Immunohistochemistry, Immunofluorescence, and Morphometry

Pancreas were fixed in 3.7% formalin solution, embedded in paraffin, and cut into 5-μm sections. Morphometrical parameters (β-cell fraction, islet size, and density) were evaluated on four sampled sections per pancreas after immunohistochemistry using a mouse monoclonal anti-insulin (Sigma-Aldrich, St. Louis, MO). Secondary antibodies coupled to horseradish peroxidase or to alkaline phosphastase were obtained from Jackson ImmunoResearch (Westgrove, PA). Enzyme substrates were DAB+ (Dako). Morphometrical parameters were determined as previously described (17) and are described in detail in the <u>Supplementary Data</u>.

### RNA Analysis

Total RNA was isolated (RNeasy Mini Plus Kit; QIAGEN, Hilden, Germany) and reverse transcribed into cDNA using the SuperScript III reverse transcriptase kit per manufacturer’s instructions (Invitrogen, Carlsbad, CA). Gene expression was quantified by real-time PCR using SYBR Green Supermix (Eurogentec, Seraing, Belgium) in a QuantStudio 1 (Applied Biosystems). The mRNA levels of each specific gene product were calculated using the ddCt method and normalized to the reference gene *RPL19* expression level. Values are expressed as the fold change of the treated compared to the control condition. Primer sequences are available upon request.

### Pancreas clearing and 3D analysis pipeline

The organs were washed, fixed, immunolabeled for insulin (anti-insulin antibody 1/800; Sigma I2018) and cleared. We then carried out acquisitions on a light sheet microscope (UltraMicroscope II Light Sheet microscope, LaVision BioTec) and a 3D reconstruction with the Arivis software follow by analysis using MeshLab software for decimation and MathLab software for quantification. Detailed protocol is provide in Supplemental data (supData1).

### Statistical Analysis

All results are presented as mean ± SEM. Comparisons were performed using the Mann-Whitney *U*, and one-way or two-way ANOVA tests. *P* < 0.05 was considered significant.

## Results

### Beta-cell adaptation is observed in response to the insulin resistance induced by either genetic-lipodystrophy or the use of an insulin receptor antagonist

Mice that were specifically deleted for Seipin in adipose tissues (Adipo Seipin KO) presented a mild insulin resistance (Fig1A and B) compared to controls. Morphometrical analysis carried on pancreatic sections of Adipo Seipin KO mice (Fig1 C and D) showed an increase in the beta-cell fraction (Fig1E) with no increase in beta-cell mass (Fig1F) or mean islet size (Fig1G) but an augmented islet density (Fig1H), suggesting that beta-cell neogenesis rather than beta-cell proliferation was responsible for the increased beta-cell fraction in response to the genetically-induced insulin resistance.

**Figure 1:**
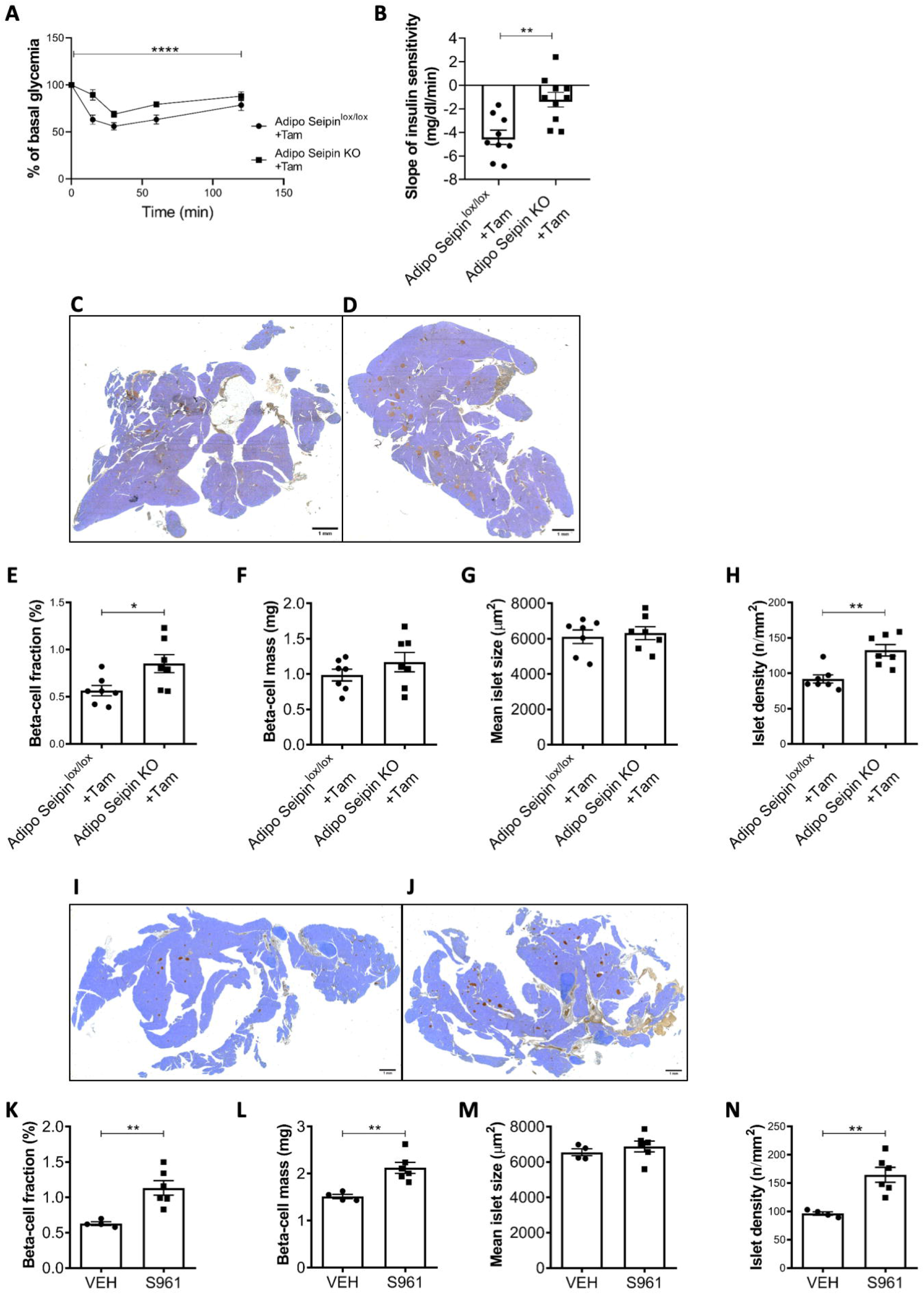
Both genetic- and pharmacological-induced insulin resistances lead to pancreatic adaptation by beta-cell neogenesis. (A) Insulin tolerance test, presented as the percentage of basal glycemia, performed on Adipo Seipin KO mice (n=9) or WT mice (n=10) and (B) slopes of early insulin sensitivity between T15 and T0. (C) and (D) representative images of insulin immunostaining (brown) of pancreatic section of WT (C) and Adipo Seipin KO (D) mice. Counterstaining was performed using hematoxylin (blue). (E-H) Pancreatic beta-cell fraction, beta-cell mass, mean islet size and islet density after insulin immunostaining of four pancreatic slides from each Adipo Seipin KO (n=7) or WT (n=7) mice. (I) and (J) representative images of insulin immunostaining of pancreatic sections of VEH (I) and S961-treated (J) mice. Counterstaining was performed using hematoxylin (blue). (K-N) Pancreatic beta-cell fraction, beta-cell mass, mean islet size and islet density after insulin immunostaining of four pancreatic slides from each S961-treated (n=6) or VEH (n=4) mice. Values are expressed as the mean ± SEM. * p<0.05; **p <0.01; ****p <0.0001 when comparing KO mice or S961-treated mice versus respective control mice.

We then studied the impact of severe insulin resistance using the insulin receptor antagonist S961 administration. Insulin resistance was strongly induced in S961-treated mice and was associated with a massive hyperglycemia after only one week of treatment, as previously described (30). Pancreatic analysis after one week of treatment revealed a doubled beta-cell fraction in S961-treated mice compared to control, as well as a 50% increase in beta-cell mass (Fig1I, J, K and L). Mean islet size remained unchanged (Fig1M) but S961 treatment led to an increase in islet density (Fig1N). Thus, severe insulin resistance with S961 led to beta-cell adaptation solely by stimulating beta-cell neogenesis with no beta-cell proliferation, suggesting again that beta-cell neogenesis was responsible for the increased beta-cell fraction in response to the S961-induced insulin resistance.

### A threshold of CORT-treatment defines functional or functional and morphometrical adaptation in young mice

Using our previously characterized model of insulin resistance (17), we searched to evaluate in young mice the doses of CORT treatment required to induce insulin resistance and analyze pancreatic adaptation. While our previous results were obtained by treating mice with 100 μg/ml CORT (C100), here we challenged eight-weeks old mice with either 25 (C25), 50 (C50) or 100 (C100) μg/ml of CORT for 4 weeks and compared to VEH-treated mice. After 4 weeks of CORT treatment, ITT showed severe insulin resistance in C100 and C50 whereas intermediate insulin resistance in C25 was observed compared to VEH-treated mice (Fig2 A). Slopes of early insulin sensitivity (Fig2 B) were higher in mice treated with C25 and C50 than in VEH-treated mice and even higher in mice treated with C100, revealing a more severe loss of sensitivity at C100. Accordingly, we measured plasma insulin levels in fed mice and observed a 6-, 17- and 15 fold increase in fasting insulin levels in C25, C50 and C100 mice compared to VEH-treated mice, respectively (Fig2 C).

**Figure 2:**
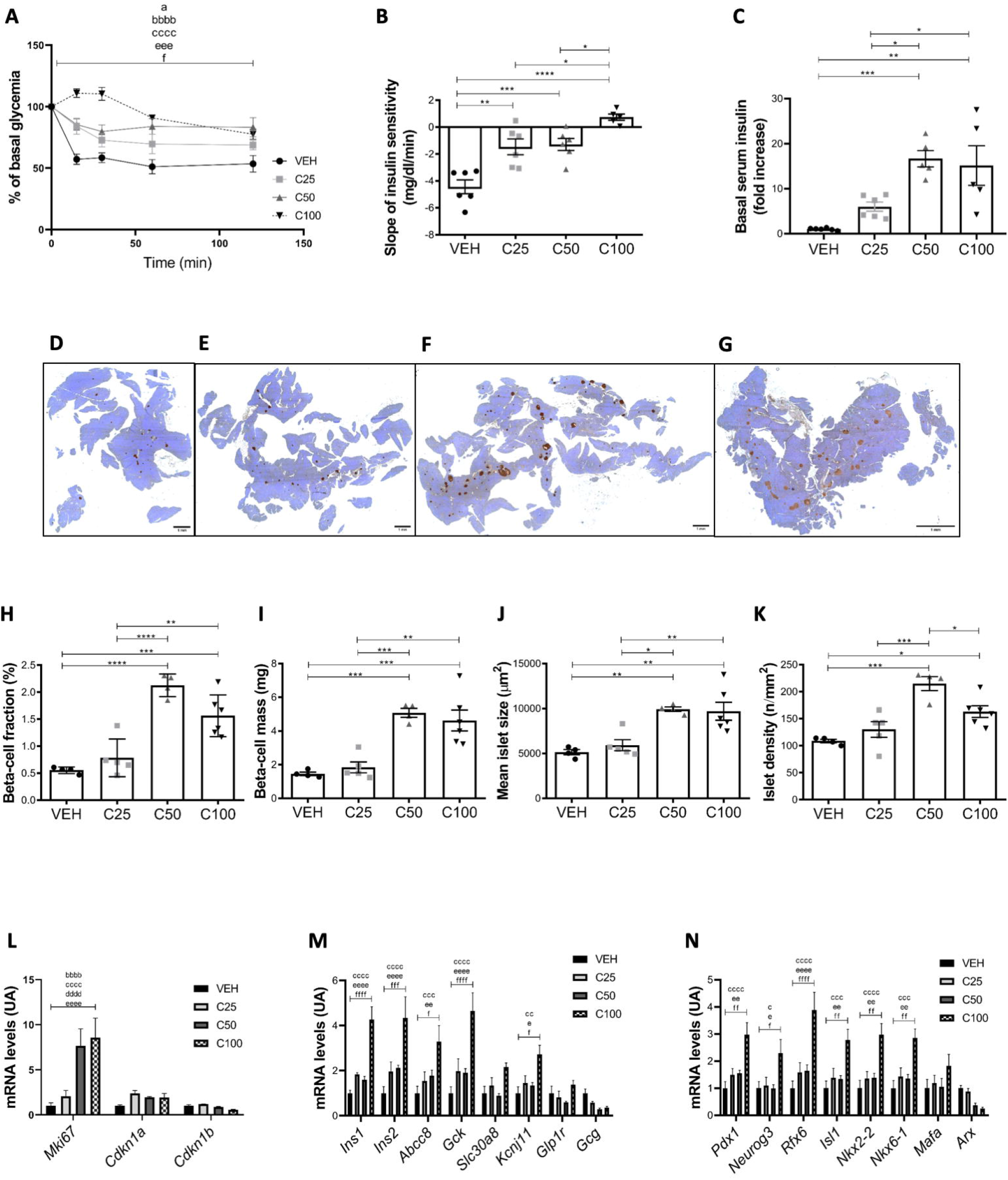
Insulin resistance severity in young mice balances beta-cell function adaptation vs function and mass adaptation. (A) ITT, presented as the percentage of basal glycemia, was carried out on mice treated with VEH (n=6), C25 (n=6), C50 (n=6) and C100 (n=5) for 4 weeks and (B) slopes of early insulin sensitivity between T15 and T0. (C) Fasted insulin serum level in mice treated with VEH (n=6), C25 (n=6), C50 (n=5) and C100 (n=5). (D-G) representative images of insulin immunostaining of pancreatic section of VEH-(D), C25-(E), C50-(F) and C100-(G) treated mice. Counterstaining was performed using hematoxylin (blue). (H-K): Pancreatic beta-cell fraction, beta-cell mass, mean islet size and islet density after insulin immunostaining of four pancreatic slides from each VEH (n=4), C25 (n=5), C50 (n=4) and C100 (n=5) mice. Values are expressed as the mean ± SEM. * p<0.05; **p <0.01; *** p<0.001; ****p <0.0001 when comparing CORT-treated versus VEH-treated mice. (L-N): mRNA levels of (L) cellular cycle, (M) beta-cell function and (N) beta-cell maturation/identity maintenance genes were determined by RT-qPCR on isolated islets from VEH-(n=6), C25-(n=5), C50-(n=5) and C100-(n=4) treated mice. Values are expressed as the mean ± SEM. a p<0.05 C25 *vs* VEH; b p<0.05 C50 *vs* VEH; c p<0.05 C100 *vs* VEH; d p<0.05 C25 *vs* C50; e p<0.05 C25 *vs* C100 and f p<0.05 C50 *vs* C100; and x<0.05; xx<0.01; xxx<0.001; xxxx<0.0001.

Morphometric analysis revealed an increase in beta-cell fraction and mass in C50- and C100-treated mice but not in C25-treated mice (Fig2H and I). Similar to C100, increase of beta-cell mass in C50-treated mice was achieved through an increase in mean islet size and islet density, suggesting both proliferation and neogenesis (Fig2 J-K). Accordingly, *Mki67* mRNA levels, a proliferation marker, were specifically increased in C50 and C100 conditions (Fig2 L). We also noticed a significant increase in the mRNA levels of several genes crucial for the function (*INS1, INS2, ABCC8, GCK, KCNJ11*) (Fig2M) and maturation/identity maintenance (*PDX1, NGN3, RFX6, ISL1, NKX2-2/6-1*) (Fig2N) of beta cells in C100 mice. Notably, despite the lack of statistical significance, we observed a tendency towards the increased expression of the same function and maturation/identity maintenance genes in C25 and C50 conditions (Fig2M and N), along with a decreased mRNA levels of *GCG* and *ARX* (Fig2 M and N), respectively important for function and identity of alpha cells.

Taken together, these results demonstrate that mild insulin resistance induced by CORT 25 treatment could lead to beta-cell adaptation solely *via* a functional increase in insulin secretion with no change in the beta-cell mass, while more severe insulin resistance (CORT 50-100) induces both functional and beta-cell mass increase through proliferation and neogenesis.

### Pancreatic adaptation is maintained in older mice and induced by mild IR

Aging is a limiting process in adaptive mechanisms: for example, treatment with the GLP-1 receptor agonist, known to increase beta-cell proliferation in young mice, failed to induce the same adaptation in mice older than 8 months (17). Thus, we sought to determine whether beta-cell adaptation capabilities in response to moderate and severe CORT-induced insulin resistance could still be observed in 12-month-old mice. To prevent excessive deaths due to CORT treatment, we only performed a 3-week CORT-treatment.

Compared to young mice, one-year-old mice present the exact same profile of ITT in response to increasing doses of CORT than the young mice, with mild insulin resistance induced at C25 and severe insulin resistance at C50 and C100 compared to VEH treated mice (Fig3A and B). Compared to VEH-treated mice, blood insulin levels in fed mice were increased by 1.8-, 7.3- and 25.9-fold in C25, C50 and C100 treated aged-mice respectively, demonstrating a maintenance of beta-cell functional adaptation in response to insulin resistance induced by CORT treatment in aged mice. Of note, compared to young mice where CORT C25 only triggered a functional adaptation of beta cells, C25 treatment in older mice resulted in an increase in beta-cell fraction and mass (Fig3D, E, H-J). A similar increase of beta-cell mass and fraction was observed at C25, C50 and C100 compared to control (Fig3 H-I). Interestingly, in C25-treated aged mice, the increase in beta-cell mass correlated with a higher mean islet size without significant change in islet density, suggesting adaptation by proliferation only, while in C50- and C100-treated mice, a significant increase in islet density was observed, indicating beta-cell neogenesis. This suggests that in extreme conditions of insulin resistance, beta-cell neogenesis associates with proliferation to allow adaptation in aged mice. Accordingly, we observed an increased expression of the proliferation marker *Mki67* in C100 mice (Fig3L). However, no significant change of beta-cell function or maturation/identity maintenance gene expression was observed in aged CORT-treated mice (Fig3M and N), except for a significant decreased expression of *GLP1R* in C50- and C100-treated mice compared to VEH-treated mice. Interestingly, we observed a decrease in mRNA levels of *GCG* and *ARX* in C50- and C100-treated mice (Fig3 M-N), similarly to young mice, suggesting a repression of alpha-cell function.

Overall, these results show that in response to severe insulin resistance induced by CORT the adaptation of beta-cell mass and function through proliferation and neogenesis is maintained in older mice.

**Figure 3:**
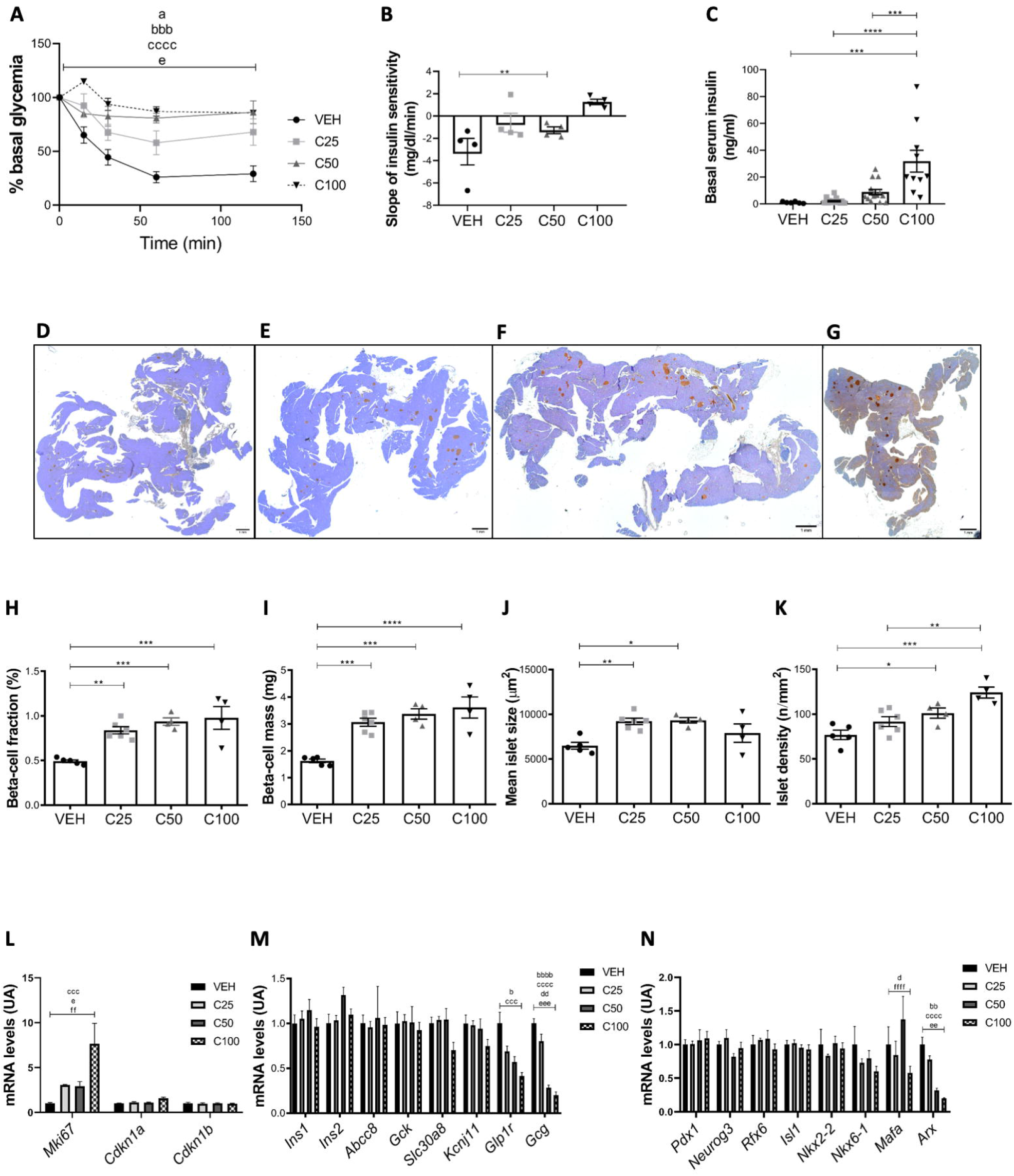
Pancreatic beta-cell adaptation is present in response to CORT-induced insulin resistance and involves both beta-cell proliferation and neogenesis in aged mice. (A) ITT, presented as the percentage of basal glycemia, was carried out on mice treated with VEH (n=4), C25 (n=4), C50 (n=4) and C100 (n=4) and (B) slopes of early insulin sensitivity between T15 and T0. (C) Fasted insulin serum level in mice treated with VEH (n=6), C25 (n=17), C50 (n=15) and C100 (n=10). (D-G) Representative images of insulin immunostaining of pancreatic section of VEH-(D), C25-(E), C50-(F) and C100-(G) treated mice. Counterstaining was performed using hematoxylin (blue). (H-K): Pancreatic beta-cell fraction, beta-cell mass, mean islet size and islet density after insulin immunostaining of four pancreatic slides from each VEH (n=5), C25 (n=6), C50 (n=4) and C100 (n=4). Values are expressed as the mean ± SEM. * p<0.05; **p <0.01; *** p<0.001; ****p <0.0001 when comparing CORT-treated versus VEH-treated mice. (L-N) mRNA levels of (L) cellular cycle, (M) beta-cell function and (N) beta-cell maturation/identity maintenance genes were determined by RT-qPCR on isolated islets from VEH-(n=4), C25-(n=4), C50-(n=4) and C100-(n=7) treated mice. Values are expressed as the mean ± SEM. a p<0.05 C25 *vs* VEH; b p<0.05 C50 *vs* VEH; c p<0.05 C100 *vs* VEH; d p<0.05 C25 *vs* C50; e p<0.05 C25 *vs* C100 and f p<0.05 C50 *vs* C100; and x<0.05; xx<0.01; xxx<0.001; xxxx<0.0001.

### Detailed 3D analysis validates pancreatic adaptation through increased beta-cell mass, islet size and density in young mice in response to CORT-induced insulin resistance

To study neogenesis in the pancreas, morphometrical techniques used until today mainly involved 2D analysis of pancreatic sections (17) or 3D analysis restricted to specific areas of the pancreas (24, 33). As a result, several biases as the cutting plan or sampling of histological slide/3D area preclude a precise counting of pancreatic islets that is necessary to fully characterize beta-cell neogenesis, as it translates in increased numbers of isolated islets, notably small beta-cell clusters. To fill this technical gap and fully validate the existence of beta-cell neogenesis in our model we developed a 3D model allowing insulin staining and tissue clearing of the whole pancreas. Based on our previous results obtain by 2D morphometrical analysis (Fig2 D-K), we chose to study C50- and VEH-treated mice, as C50 mice show a strong and homogenous adaptation in young adult mice. Our protocol allowed for insulin-staining and tissue clearing of whole pancreas, to *in fine* assess critical parameters such as whole organ volume, whole insulin-immunofluorescence volume, counting and classification by individual volume of insulin-positive cell clusters (Fig4).

**Figure 4:**
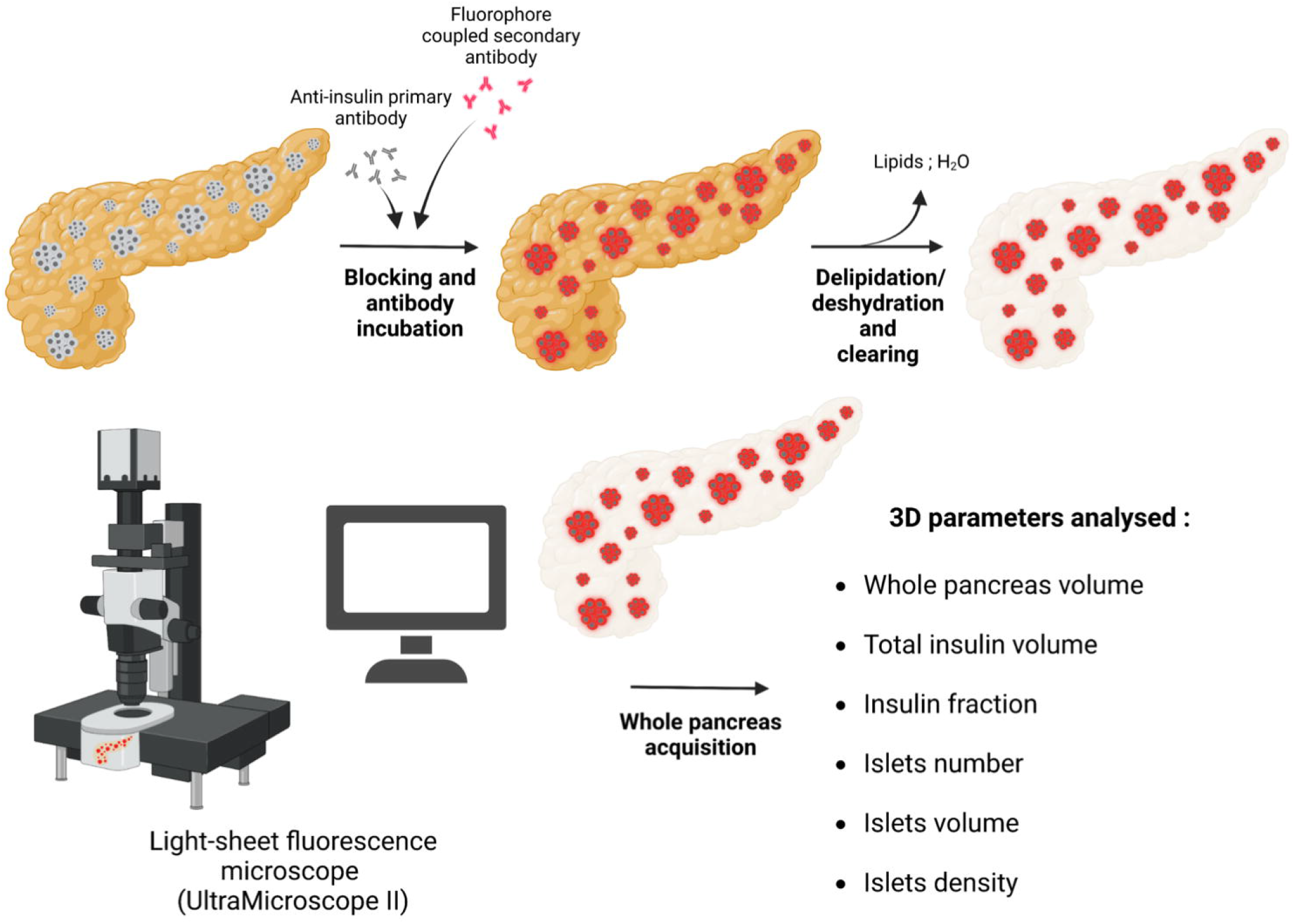
Schematic representation of the three-dimensional analysis protocol. Pancreas were isolated after sacrifice of 3-month or 12-month-old, VEH or C50-treated mice. After blocking, pancreases were sequentially incubated with primary and secondary antibodies. Stained-pancreases then undergo dehydrative and delipidant washes before clearing by correction of refractive index. Acquisitions on whole pancreases were made using a light-sheet fluorescence microscope. 3D reconstruction then allowed analysis of morphometrical parameters such as total pancreas volume (autofluorescence), total insulin volume (fluorescent insulin staining) and other parameters obtained with a sufficient resolution to discriminate islets individually.

A short summary of the protocol is provided in figure 4. Briefly, whole pancreas were fixed, incubated with primary antibodies and secondary antibodies. Then, pancreases were dehydrated, delipidated and cleared with organic solvants. Finally, Z-stacks acquisition of whole pancreas were performed on a light-sheet microscope at 2.8 magnificent with a z-step of 1 or 10μm for insulin staining or volume acquisition, respectively. A complete protocol for the procedure is provided as supplementary material (supData1).

First, visualization of Z-stacks obtained after whole pancreas acquisition of both C50- and VEH-treated animals confirmed a specific and homogenous staining of insulin-positive cells, regrouped in well-defined circular objects, classified as pancreatic islets (Fig5 A-B). Following objects/islets’ segmentations (supData1), three-dimensional reconstructions of whole pancreas show a regular distribution of islets throughout the pancreas, validating our pipeline for image segmentation (Fig5C and D).

**Figure 5:**
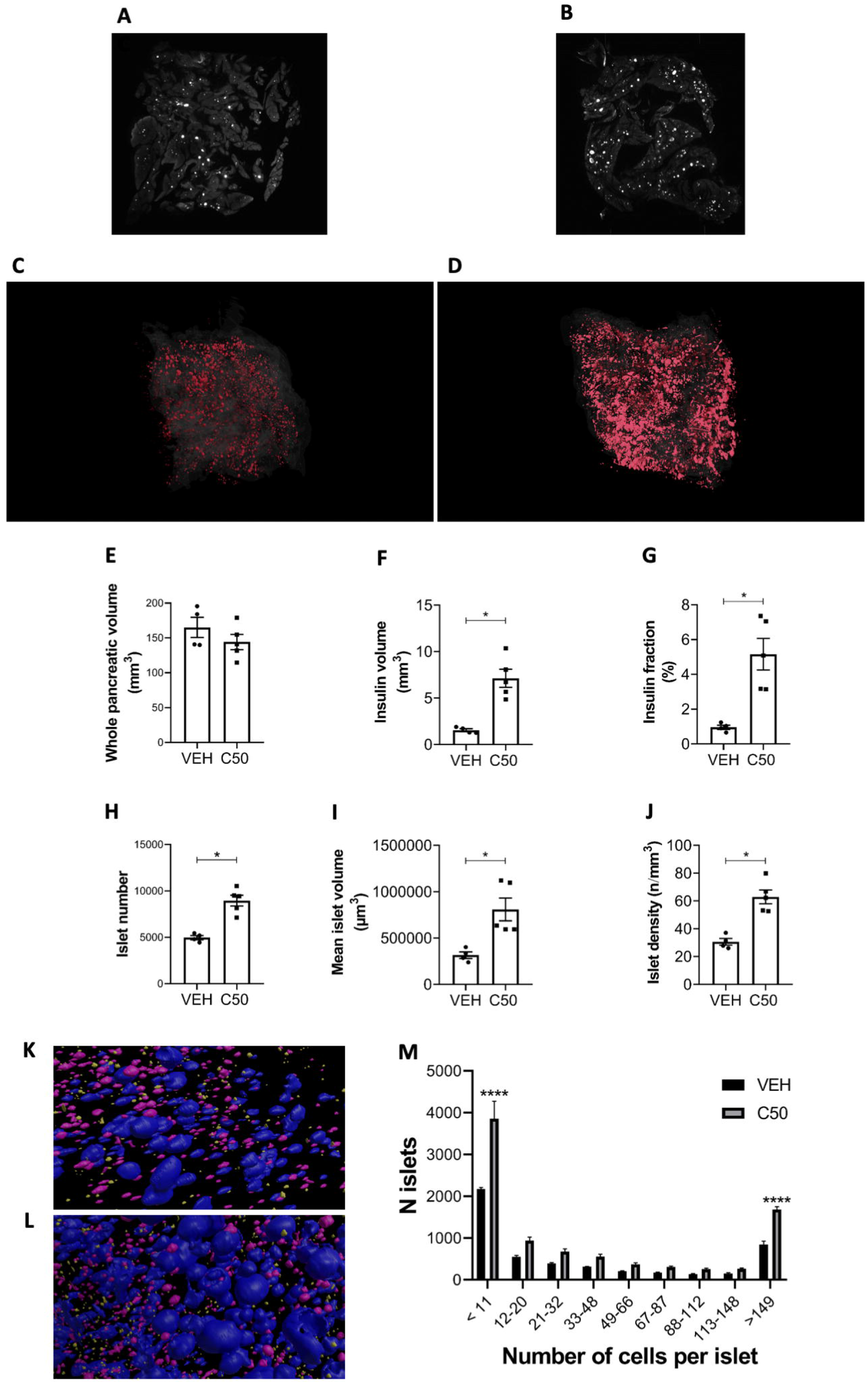
3D analysis of pancreas from CORT-induced insulin resistant mice confirms increased islet numbers as an adaptive mechanism to increase beta-cell mass in young adult mice. (A-B) Representative images of a Z-stack section after acquisition on whole pancreas stained for insulin using light-sheet microscopy from VEH (A) and C50 (B) mice. (C-D) representative images of the 3D reconstruction following image segmentation of volume (glass-like structure) and islets (red structures) on whole pancreas from VEH (A) and C50 (B) mice. (E-J) Whole pancreas volume; total insulin volume; insulin fraction; islet number, mean islet volume and islet density calculated after image segmentation and analysis of pancreas from VEH (n=4) and C50 (n=5) mice. (K-L) Illustration of islet size diversity and density inside a pancreas of VEH (K) and C50 mouse (L). (M) Islet size distribution in pancreas from VEH (n=4) and C50 (n=5) mice. Size categories were defined as indicated in Supplemental data. Values are expressed as the mean ± SEM. * p<0,05 when comparing CORT-treated versus VEH-treated mice.

In agreement with our previous findings (17), whole pancreatic volume was identical in VEH and C50 mice (Fig5 E) and whole insulin volume, obtained by summing the individual volume of each segmented islet, was 4.6 times higher in C50 mice compared to VEH (Fig5F). Similarly, we measured a 5.4fold increase of insulin fraction in C50 mice compared to VEH (Fig5G). Islet counting showed a mean number of 8956.2 +/-1324 *vs* 4981 +/-440.6 islets per pancreas in C50-treated mice compared to VEH-treated mice, indicating 80% more islets in C50 compared to VEH mice (Fig5H; supMovie1A-B), suggesting beta-cell neogenesis. Comparison of individual mean islet volume showed a 2.6-fold higher volume in C50 mice compared to VEH mice (Fig5 I), suggesting increased proliferation of beta cells. By relating the number of islets to the whole pancreas volume, we demonstrated a 2-fold increase in islet density, i.e. 63+/-11.21 SD *vs* 30 +/-4.91 SD islets/mm^3^ of pancreatic tissue in C50 compared to VEH mice (Fig5J). To further characterize islets’ neogenesis and proliferation, we performed islet classification, as previously used for 2D analysis (17): islets were sorted in 9 classes of beta-cell clusters based on volume analysis starting from small islets cluster (i.e. containing 11 cells or less) to larger islets with more than 149 cells (classes are described in supData1). We observed a statistical almost 2-fold increase in both the smallest and the biggest classes of cell clusters (Fig5K ; supMovie2A-B). Although it did not reach statistical significance, we could also observe a trend towards an increase in the number of islets of every other classes, with an average factor of 1.7 fold increase (Fig5 K).

Taken together, these results unequivocally show increased beta-cell mass and fraction up to a 5-fold in CORT-treated mice. They also demonstrate the implication of proliferation in pancreatic adaptation to insulin resistance, with a doubled mean islet volume. Importantly, our results offer a solid confirmation of the presence of beta-cell neogenesis in our young adult mouse model of severe insulin resistance, with a doubled islet density and a clear and specific raise in small islets number. Using a whole-organ analysis approach, we demonstrate without spatial or structural biases the equal involvement of both proliferation and beta-cell neogenesis in an adaptive process in young adult mice.

## Discussion

In our study, we demonstrated that insulin resistance is a princeps signal triggering pancreatic adaptation through both a functional increase in insulin secretion and/or a beta-cell mass increase *via* either beta-cell proliferation and neogenesis. Importantly, we demonstrated that pancreatic adaptation by beta-cell neogenesis is a conserved mechanism in both mild and severe insulin resistance and in young and older mice, establishing beta-cell neogenesis as an important adaptive mechanism that could be therapeutically triggered in the future to endogenously compensate for insulin deficiency in adult. Strikingly, we developed an unique 3D light sheet immunofluorescence analysis of insulin positive cells to demonstrate that GC-induced insulin resistance is associated with an increase in both islets’ size and number reflecting proliferation and neogenesis, respectively.

While adult beta-cell neogenesis was suggested but not quantified in models of pancreatic injury in mice, studies failed to confirm its implication, thereby remaining a controversial adaptive mechanism (34). Conversely, some results proposed that the major contributor to increase and/or maintain the beta-cell mass in adult rodents is beta-cell replication (35, 36). Previously, we demonstrated, by 2D analysis techniques on sampled pancreas sections, an increase in the density of islets, notably small neo formed islets (17). Yet, biases associated with the 2D analysis prompted us to develop organ clearing that represents an advanced technology for a detailed morphometrical analysis of entire pancreas. Recently, a study using macroscopic 3D analysis on pancreas from S961-treated, diet-induced obese mice, has reported increased numbers of islets mostly attributed to beta-cell proliferation (37). Moreover, authors conducting 3D analysis by optical projection tomography on ob/ob mice revealed no augmentation in the number of small islets (that could reflect the presence of beta-cell neogenesis) but acknowledge beta-cell hyperplasia as the main adaptive mechanism in their model (38). However, whole 3D analysis of the endocrine pancreas with a particular focus on islets neogenesis was never performed on a model of severely induced-insulin resistance. To fill this caveat, we adapted a protocol to measure beta-cell and islet populations in whole pancreases. Our sensitive insulin staining enabled us to identify even small beta-cell clusters containing as few as 1-3 beta cells. We showed in adult young mice a clear and unbiased increase in beta-cell mass and fraction with a doubled islet number following insulin resistance, and confirming our 2D analysis. A more detailed analysis of islet populations showed an increase in both small and large beta-cell cluster, thereby evidencing that beta-cell neogenesis, and beta-cell proliferation are both essential mechanisms to pancreatic adaptation in adult young mice.

However, the origin of the neo-formed beta cells remains unknown. Several studies in fact proposed a ductal origin of endocrine precursors and highlighted the importance of master transcription factors such as HNF1B and NGN3 in adult beta-cell neogenesis. (39, 40). A recent lineage tracing study mixed with single cell RNAseq and microscopic 3D imaging showed the implication of the endocrine master transcription factor NGN3 in beta-cell homeostasis in adult and suggested that ductal NGN3 was involved in a mouse model of type 1 diabetes (33). To gain further insight into the role of duct in beta-cell neogenesis, we propose that whole pancreas 3D analysis of both ducts and islets could allow the study of critical parameters such as ductal network density, ducts-to-islets distance measurement and the dynamic evolution of duct and islets networks in response to proneogenic stimuli. *In fine*, our experimental settings allowed us to take a snapshot of the 3D structure of pancreatic islets and ducts, suggesting a close proximity in-between islets and ducts (data not shown), however further evidences and analysis are needed to demonstrate the ductal origin of neo-formed islets.

Lipodystrophic model of insulin resistance (Bscl2^lox/lox^ × Adipoq-CreER mice) display pancreatic adaptation preferentially by neogenesis despite mild insulin resistance but again proposing that insulin resistance could be the driving event for beta-cell adaptation through beta-cell neogenesis. Interestingly, mice that were heterogously deleted for the insulin receptor and IRS-1 displayed severe insulin resistance associated with an increase in beta-cell mass. However, islets counting was not performed on this model even though the presented data suggested that beta-cell neogenesis was indeed present (41). Nevertheless, a murine model presenting a monogenic form of mild insulin resistance through global IRS1 knockout showed what authors called a spotty insulin staining observed throughout the exocrine pancreas as well as an increased proportion of small islets compared to WT mice strongly suggesting beta-cell neogenesis (42). Thus, beta-cell neogenesis could be observed in several other forms of genetically-induced insulin resistance. Moreover, while pharmacological insulin resistance with the insulin receptor antagonist S961 has been shown to be associated with increased beta-cell proliferation without evidence of beta-cell neogenesis (24, 43). Here we demonstrated that the severe insulin resistance induced by a short-term S961 treatment stimulates a rapid and exclusive adaptation through islets neogenesis. The unaltered mean islet size of S961-treated mice, a classical landmark of beta-cell proliferation, suggests that the first mechanism implemented to rapidly meet the body’s insulin needs is the formation of new beta cells and not the proliferation of preexisting cells. Thus, the severity of S961-induced insulin resistance may explain this dichotomy in pancreatic adaptation mechanisms.

To further assess the link between the degree of insulin resistance and level of pancreatic adaptation, we treated young and older mice with different doses of CORT to achieve gradual levels of insulin resistance. We demonstrated that in young mice mild CORT-induced insulin resistance leads to beta-cell adaptation solely by increasing secretory capacity without stimulating mass adaptation. More severe insulin resistance however did trigger both function and mass adaptation *via* proliferation and neogenesis. Similarly to humans, middle- to old-aged mice naturally develop insulin resistance and can compensate for the increase in glycemia by a higher insulin secretion, suggesting a pancreatic adaptation that likely does not result from an increase in pre-existing beta-cell proliferation since proliferation rate of beta cells remains low at adult stage in Human and mice (0,1 – 0,3% a day) (44-46). Interestingly, in our study, older mice exposed to CORT treatment become insulin resistant and are still able to recruit beta-cell function and mass in order to increase insulin production. Contrary to young mince, beta-cell adaptation even seems potentialized at the lowest dose of CORT (C25) that was sufficient to trigger a beta-cell mass increase. This suggests that beta-cell functional adaptation, while preserved, is no longer sufficient to compensate insulin resistance induced by age and C25 treatment and requires beta-cell mass augmentation. Finally, if beta-cell proliferation is present in all groups of aged CORT-treated mice, it reaches a plateau at C25 dose and beta-cell neogenesis is then stimulated at C50 and C100 doses. Other protocols to induce insulin resistance such as a 1 week to 12 months high fat diet are known to increase beta-cell mass by proliferation only (47, 48) and did not require explicit insulin resistance to start the pancreatic adaptive program. In contrast, short term daily injections with GC at various doses (dexamethasone) led to gradual insulin resistance and an increase in basal insulin secretion partly explained by beta-cell mass expanse through proliferation. But then again, beta-cell neogenesis was not fully investigated in these models (49-51).

Taken together, our results offer a broad view of beta-cell neogenesis as an adaptive mechanism in several situations of insulin resistance, either genetic or drug-induced. Our study demonstrates that the level of pancreatic adaptation depends on both duration and severity of insulin resistance. We defined that age is not a barrier to pancreatic adaptation and that beta-cell mass increase *via* neogenesis compensates for an age-related decline in beta-cell function to ensure sufficient insulin production. The use of an innovative, whole organ 3D analysis allowed us to show a massive increase in beta-cell mass through neogenesis and could be further used as a tool to monitor the efficacy of therapeutic strategies aiming at increasing beta-cell mass. Beta-cell neogenesis could be the best strategy to cure type 1 diabetes because the nearly total absence of beta cells in the pancreas of T1D patients precludes from developing regenerative strategies relying on pre-existing beta cells only. Being able to stimulate beta-cell precursors differentiation in adult has been demonstrated in our previous and present study but translating this result into therapy remains a challenge. As previously described (17), serum from CORT-treated mice contains factors that can stimulate beta-cell neogenesis. The origin of the neogenic factors remains to be defined, even though we hypothesized a peripheric origin of the factors coming from insulin sensitive tissues. Complex strategies using both transcriptomic, secretomic of tissues rendered insulin resistant need to be developed to identify circulating factors that induce pancreatic beta-cell adaptation, with a specific focus on factors able to stimulate beta-cell neogenesis and/or improved insulin secretion by beta cells.

## Acknowledgments

This work was funded by INSERM, Sorbonne Université, Fondation pour la Recherche Médicale (FRM EQU201903007868), Société Francophone du Diabète, Aide aux Jeunes Diabétiques, Type 1 running Team and World Diabetes Tour. We gratefully aknowledge the UtechS Photonic BioImaging (Imagopole), especially Julien Fernandes, C2RT, Institut Pasteur, supported by the French National Research Agency (France BioImaging; ANR-10– INBS–04; Investments for the Future). The authors thank T. Ledent, L. Dinard, A. Guyomard, T. Coulais, and Q. Pointout (animal housing facility, Saint-Antoine Research Center, Sorbonne University, INSERM, Paris). A.L. was supported by doctoral fellowship from Ministère de l’Enseignement Supérieur et de la Recherche

## Supplemental data

### Supplemental data 1 : Detailed three-dimensional analysis protocol

Mice were anesthetized using isoflurane and received through intracardiac infusion 10ml of cold PBS and 20ml of cold 4% PFA. The dissected pancreases were stored in 20ml of 4% PFA at 4°C for a maximum period of one month. Pancreas staining and clearing is achieved following iDisco clearing technique with an adapted timing. Pancreases were incubated three times with PBS +0.2% Gelatin (Sigma ref 48723-500-F) + 0.5%TRITON 100X (SigmaT8787) (PBSGT) during one hour at 37°C and incubated in PBSGT four 3 days at 37°C under slow agitation to block nonspecific binding and permeabilize the pancreases.

Permeabilized pancreases were incubated in PBSGT with 0,1-0,2% saponin (Sigma ref S4521) (PBSGT-sap) and mouse anti-insulin antibody (1/800; Sigma I2018) during 5 days at 37°C under slow rotation. Pancreases were washed 6 times in PBSGT at 37°C during 1h under slow agitation. Pancreases were incubated in PBSGT-sap with Alexa 750 coupled anti-mouse antibody (1/50; Life Technologies A21037) for at least 3 days at 37°C under slow agitation. Pancreases were washed 6 times in PBSGT at 37°C during 1h under slow agitation. At this step, pancreas were either stored in PBSGT at 37°C without agitation for 1 day or directly dehydrated, as follows.

For dehydration, pancreases were first washed two times in H2O MilliQ at room temperature during 1h under slow agitation, and then, incubated at room temperature under slow agitation in methanol solutions (20 %, 40%, 60%, 80% methanol diluted in H2O MilliQ and two times in 100% methanol) during 1h for each bath. Pancreases were then incubated over-night in 1/3 methanol + 2/3 dichloromethane (DCM; Sigma ref 270997) solution. Complete delipidation was achieved by 2 incubations in pure DCM bath at room temperature during 1h each incubation under slow agitation. Right after delipidation, pancreases were incubated in dibenzyl ether (DBE; SIGMA ref 33630) to allow refractive index correction.

Acquisitions of whole pancreas were performed in collaboration with Pasteur Institute using UltraMicroscope II Light Sheet microscope. Volume acquisition was performed using tissue autofluorescence (470/40nm excitation filter; 525/50 nm emission filter) with a z-step of 10μm; islets acquisitions were performed with a z-step of 1μm (710/75nm excitation filter; 785/62nm emission filter), both using horizontal dynamic focusing.

Pancreatic islets were segmented automatically using an analysis pipeline in Arivis Vision 4D (v3.1.1, Arivis AG, Rostock, Germany). Briefly: The islet channel in the 3D image was first filtered using a 2D median of radius of 3 pixels. Then two different steps of segmentation were performed. First islets were segmented on the median-filtered image using a threshold on intensity. The value of this threshold was adapted for each image, depending on the preservation of the markers, and range from 180 to 1800. Spurious objects were filtered out by rejecting objects with a 3D sphericity (without holes) below 0.3, and objects smaller than 100 voxels. The resulting 3D binary mask was then smoothed using the plane-wise ‘Closing’ morphological filter with a radius of 3 pixels. Because this filtering discarded big islets that had a complex shape and hence, a low sphericity, we used an additional segmentation to rescue them. A second identical step of segmentation was performed, on the same filtered image, but with thresholds on intensity about 50% larger than in the first step and a threshold on sphericity of 0.1. The two binary 3D masks were then exported as meshes, and post-processed in MeshLab (52) (v2022.2).

In MeshLab, the islets meshes were decimated using the Quadratic Edge Collapse Decimation method, reducing the number of vertices from 40M to about 5M per image.

The two meshes from the two segmentation steps were then combined using MATLAB (R2021a, The Matworks, Natick, USA) to produce a single mesh file where each islet is represented by one closed mesh. These individual meshes were then used to yield estimate of the volume, intensity etc.

The pancreas volume was segmented from the autofluorescence signal, on an image with larger pixel size, using again Arivis Vision4D. The image was first filtered with a 2D median of radius 3 pixels, then segmented with a threshold on intensity above 200. The resulting 3D binary mask was filtered with the ‘Closing’ morphological plane-wise operation with a radius 5 pixels.

Islets classification was determine based on our previous method (17) by calculating the volume of an individual beta cell (≈1670 μm3) to obtain approximative volume of beta-cell clusters/islets as following :

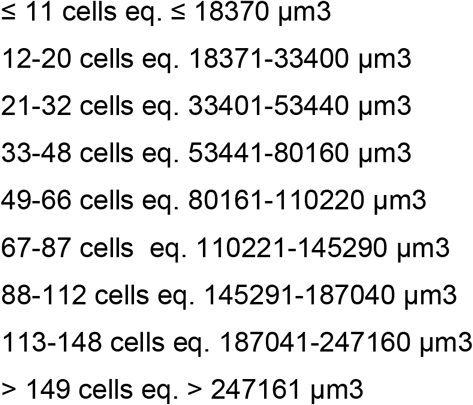

Reconstructions presented as Figure 5 C-D and films in Supplemental data (supMovie1-2) were performed on Blender software using meshes post-process as described above.

